# Clinicopathologic Dissociation: Robust Lafora Body Accumulation in Malin KO Mice Without Observable Changes in Home-cage Behavior

**DOI:** 10.1101/2023.09.11.557226

**Authors:** Vaishnav Krishnan, Jun Wu, Arindam Ghosh Mazumder, Jessica L. Kamen, Catharina Schirmer, Nandani Adhyapak, John Samuel Bass, Samuel C. Lee, Atul Maheshwari, Gemma Molinaro, Jay R. Gibson, Kimberly M. Huber, Berge A Minassian

## Abstract

Lafora Disease (LD) is a syndrome of progressive myoclonic epilepsy and cumulative neurocognitive deterioration caused by recessively inherited genetic lesions of EPM2A (laforin) or NHLRC1 (malin). Neuropsychiatric symptomatology in LD is thought to be directly downstream of neuronal and astrocytic polyglucosan aggregates, termed Lafora bodies (LBs), which faithfully accumulate in an age-dependent manner in all mouse models of LD. In this study, we applied home-cage monitoring to examine the extent of neurobehavioral deterioration in a model of malin-deficient LD, as a means to identify robust preclinical endpoints that may guide the selection of novel genetic treatments. At 6 weeks, ∼6-7 months and ∼12 months of age, malin deficient mice (“KO”) and wild type (WT) littermates underwent a standardized home-cage behavioral assessment designed to non-obtrusively appraise features of rest/arousal, consumptive behaviors, risk aversion and voluntary wheel-running. At all timepoints, and over a range of metrics that we report transparently, WT and KO mice were essentially indistinguishable. In contrast, within WT mice compared across timepoints, we identified age-related nocturnal hypoactivity, diminished sucrose preference and reduced wheel-running. Neuropathological examinations in subsets of the same mice revealed expected age dependent LB accumulation, gliosis and microglial activation in cortical and subcortical brain regions. At 12 months of age, despite the burden of neocortical LBs, we did not identify spontaneous seizures during an electroencephalographic (EEG) survey, and KO and WT mice exhibited similar spectral EEG features. Using an *in vitro* assay of neocortical function, paroxysmal increases in network activity (UP states) in KO slices were more prolonged at 3 and 6 months of age, but were similar to WT at 12 months. KO mice displayed a distinct response to pentylenetetrazole, with a greater incidence of clonic seizures and a more pronounced post-ictal suppression of movement, feeding and drinking behavior. Together, these results highlight a stark clinicopathologic dissociation in a mouse model of LD, where LBs accrue substantially without clinically meaningful changes in overall wellbeing. Our findings allude to a delay between LB accumulation and neurobehavioral decline: one that may provide a window for treatment, and whose precise duration may be difficult to ascertain within the typical lifespan of a laboratory mouse.

## Introduction

Lafora disease (LD) is a recessively inherited disorder of glycogen metabolism that gives rise to a syndrome of adolescent-onset progressive neurocognitive decline and intractable myoclonic epilepsy^1-5^. LD is pathologically defined by the buildup of Lafora bodies (LBs): intracellular periodic acid-Schiff (PAS)-positive inclusions of polyglucosan^6^ that are found in neurons and astrocytes, as well as muscle, liver and skin^1,7^. LD is caused by mutations in either of two genes located on chromosome 6: *EPM2A* (encoding laforin, a carbohydrate binding dual-specificity phosphatase^8^) or *NHLRC1* (encoding malin, an E3-ubiquitin ligase^9^). Current working models of pathophysiology hypothesize that malin and laforin function as a quality control complex within growing glycogen polymers, preventing the generation of very long branches that promote precipitation^1,2,10,11^. A deeper knowledge of these mechanisms remains an active area of research, focusing on a detailed understanding of the substrates of laforin and malin’s phosphatase and E3-ligase activities, respectively. In the absence of either malin or laforin, insoluble aggregates of glycogen are hypothesized to trigger a cascade of neuroinflammation, neuronal dysfunction and hyperexcitability. Despite our fairly advanced genetic understanding of LD, today’s standard of care remains supportive, with death commonly occurring within 10 years of initial diagnosis, typically due to status epilepticus or aspiration pneumonia^2,12^.

Multiple LD mouse models have been developed through targeted constitutive deletions of either *EPM2A*^6,13^ and *NHLRC1*^*14-17*^. Laforin- and malin knockout (KO) mice generated in this fashion faithfully display age- and tissue-dependent accumulation of LBs, astrogliosis and microglial activation^6,17,18^, complementing their construct validity with robust pathological validity. However, evidence for face/symptomatic validity remains considerably less robust, as neither malin nor laforin KO mice display progressive *and/or* disabling myoclonic seizures, obvious cumulative frailty, or reproductive/survival deficits^19^. In one study, approximately one-year old malin KO mice displayed mild open field hypoactivity, subtle rotarod abnormalities and poorer object recognition scores^14^, together taken to reflect motor and memory deterioration. With video-electroencephalopgraphy (EEG), both myoclonic jerks and spike/wave discharges were observed in KO mice, but these often occurred independently/asynchronously. KO mice displayed an increased sensitivity to the chemoconvulsant pentylenetetrazole (PTZ), resulting in more frequent and rapidly occurring myoclonic and generalized seizures^20,21^. However, neither study compared KO mice to littermate controls^6,14,20^. A contemporaneously (but independently) generated line of malin KO mice was found to display open field *hyper*activity (interpreted as “reduced anxiety”), together with *enhanced* long-term potentiation (LTP) and unchanged rates of operant learning^17^. These KO mice displayed more pronounced and frequent clonic seizures following kainic acid injections but were not surveyed for spontaneous seizures. A third separately derived line of malin KO mice^16^ was not tested on the open field, but was found to display elevated rates of context-dependent freezing responses^22^, potentially in accordance with enhanced LTP rates. Through video observations alone, these mice were found to have elevated rates of spontaneous myoclonus (∼1.5 jerks/min, compared to 0.5 in WT littermates). And finally, an independent fourth line of KO mice^15,19^ has been extensively utilized for biochemical studies^23-25^ but has never been examined for neurobehavioral or epileptic abnormalities.

A wide range of exciting targeted genetic^26,27^ or biochemical^19,28^ treatment strategies for LD are on the horizon. To rapidly test and improve upon these therapeutics in the preclinical domain, detailed histopathological endpoints in LD genetic models could ideally be complemented by rigorously obtained endpoints of neurobehavioral decline. To this end, we sought to identify robust evidence for progressive neurobehavioral deterioration in malin KO mice^16^ using home-cage monitoring (HCM), a technique that continues to gain popularity in behavioral neuroscience. As an alternative to traditional “out of cage” assays (e.g., open field/elevated plus maze tests), HCM platforms emphasize the automated and experimenter-free collection of *prolonged* behavioral recordings that encompass the nocturnal period^29^, obtained in an enclosure adopted by the mouse as *its* home-cage. Advances in videotracking and home-cage instrumentation now permit the simultaneous assessment of multiple streams of scalar variables (e.g., feeding *and* wheel-running *and* licking, etc), allowing investigators to appraise spontaneous behavior across a variety of time scales^30,31^. Programmed provocative maneuvers/stressors, applied *within* the home-cage, further mitigate the observer effects associated with human exposure^32,33^. HCM has been applied in this manner to uncover phenotypes in mouse models of pervasive neurodevelopment^34-37^, aging^38^ and neurodegeneration^39-41^. Here, we conduct a home-cage behavioral assessment of malin KO mice and their wildtype littermates at 6 weeks, ∼6 months and 1 year of age. At this final timepoint, we survey WT and KO mice for spontaneous seizures, and apply home-cage monitoring to carefully examine the acute and subacute behavioral responses to PTZ. At each stage, we qualitatively assess markers of microglial activation, astrogliosis and LB accumulation, and quantitatively appraise neocortical circuit dysfunction through in vitro recordings of bursting behavior^42^.

## Methods

### Mice

All protocols were approved by the UTSouthwestern Medical Center (UTSW) and Baylor College of Medicine (BCM) Institutional Animal Care and Use Committees and conducted in accordance with USPHS Policy on Humane Care and Use of Laboratory Animals. 7 wildtype (WT) and 9 KO^16^ were shipped from UTSW to BCM, from which heterozygous breeding pairs were generated (perpetuating their existing genetic background). PCR-based genotyping was performed on tail DNA at ∼p16, and mice were weaned into gender-matched cages at p21. No mice were excluded.

### Home-cage Monitoring

Cohorts of age-matched WT and KO mice were transferred from the vivarium to Noldus Phenotyper home-cages (30×30×47cm) within a designated satellite study area^34,35^. The “6 week” cohort were 6.08 + 0.14 weeks of age [mean + standard deviation]), “6 months” were 28.77 + 1.95 weeks of age (delayed due to COVID restrictions on satellite use) and mice in the “1 year” cohort were 50.75 + 1.3 weeks of age. Sample sizes for each genotype are provided within figures, and a total of 13 WT mice and 16 KO mice were examined serially at each timepoint. 16 home-cages were employed in groups of four (“quad units”). Each cage contained (i) two lickometered water sources (0.8% sucrose-drinking water Vs drinking water), (ii) an infrared (IR)-lucent shelter, an aerial IR camera and IR bulb arrays, (iii) a beam-break device to measure entries into a food hopper, (iv) a detachable running wheel, and (v) a 2300Hz pure tone generator and an LED house light. Satellite temperature (20-26C), humidity (40-70%) and light cycle settings (ON between 0500-1700) matched vivarium conditions. White noise was played continuously, and satellite access was restricted to gowned, gloved, face-masked and capped personnel to minimize olfactory variations. Mice were distributed randomly to home-cages ensuring that one gender or one genotype was not over-represented within a single quad unit. When conducting within-cage daytime tasks described (e.g., positioning running wheel), operators were blinded to genotype.

### Data Acquisition

Live videotracking (Noldus Ethovision XT14) sampled the arena-calibrated x-y coordinates of the object centerpoint at 15Hz, providing rich location time series data, enabling heat maps and measures of sheltering, horizontal displacement and “sleep”. “Sleep” (enquoted to emphasize the noninvasive assessment) was defined as a period of continuous immobility lasting ≥40s, previously validated to provide >90% agreement with neurophysiologically-determined sleep^43-46^. A modular design^34,36^ was applied, beginning with a 2h-long initial habituation study (“Intro”) followed by two consecutive 23h long baseline recordings (1500-1400). Then, we applied visual and auditory stimulation (“light spot^34,35,47^ and beep^34^”), followed by a third prolonged recording in the presence of a running wheel (1400-1100). We concluded with a single daytime “cage swap”^35,36^ provocation, swapping each mouse into a cage inhabited by a conspecific of the same sex for 2h. Lickometer-registered epochs of fluid consumption were registered as lick bouts/occurrences and durations. Similarly, both feeding entries/occurrences and feeding entry durations were tallied.

### Pentylenetetrazole (PTZ) Injections

A separate cohort of 12-13 month old WT and KO mice were acclimated to home-cage chambers for 24h, following which they all received intraperitoneal injections of PTZ (Sigma-Aldrich, 30mg/kg) within the home-cage at ∼1200h, as previously described^35^. We collected a 3h long “ictal” recording beginning immediately following injection, followed by a more prolonged “post-ictal” recording (from 1600 to 1100 the following day). Through the integrated visualization of video and high resolution mobility data^35^, the first 20 minutes of the “ictal” recording was scored manually for the occurrence and latency of clinically evident seizure semiologies as classically described^48^, including features of phase 2 (partial clonus), phase 3 (generalized clonus with sudden loss of upright posture) and phase 4 (tonic-clonic maximal seizures, with or without hindlimb extension). Examples are provided in Supplemental Movies 1-3.

### Electroencephalography

To survey for spontaneous seizures, ∼12-13 month old WT and KO mice were implanted with EMKA easyTEL S-ETA devices under sterile precautions and isoflurane anesthesia^36^. Biopotential leads (2) were affixed epidurally in right frontal and left posterior parietal regions using dental cement, with wires tunneled to a transponder positioned in the subject’s left flank. Wireless EEG was acquired at 1000Hz sampling rate with IOX2 software (EMKA Technologies) via easyTEL receiver plates placed underneath home-cages, and EEG signals were inspected with LabChart reader using a bandpass filter (1-30Hz). Spectral analysis of unfiltered EEG (EEG ToolKit, Matlab) was conducted on randomly chosen 10-minute segments of wakefulness, collected between 1700 and 2000. Recordings were analyzed for power between 2 and 200 Hz, providing absolute power (AP) in dB (log10 (μV^2^/Hz)) at 1 Hz intervals. Relative power (RP) was calculated by dividing the absolute power for each frequency by the total power (TP), and then normalized with a log transformation before comparison between mice (RP=AP/TP), similar to methods described previously^49,50^.

### Histology

For histological assessments, mice were rendered comatose with a single intraperitoneal injection of 0.1ml Beuthanasia-D (phenytoin 50mg/ml/pentobarbital 390mg/l) and subsequently transcardially perfused with ice cold phosphate buffered saline (PBS) followed 10% neutral buffered formalin (NBF). Mice that had been exposed to PTZ or implanted with EEG devices were not utilized for histology. Brains were extracted, post-fixed for another 24 hours in NBF and then stored in a solution of 70% ethanol. Paraffin-embedded brain tissues were sectioned and stained using standard histological technique including diastase-digested periodic acid-Schiff (PASD) staining to visualize polyglucosan bodies^22^. Stained slides were scanned using a Hamamatsu Nanozoomer 2.0 HT digital slide scanner (40 x objective). For co-immunostaining, paraffin-embedded brain sections were deparaffinized and rehydrated by processing with xylene, decreasing concentrations of ethanol in water, and subjected to antigen retrieval using citrate buffer pH6.0 (C9999, Sigma-Aldrich). Sections were blocked with 5% normal donkey serum (in 0.1% Triton X-100, PBS) for 1 h and incubated for 48 h at 4°C with primary antibodies diluted in blocking solution, including those targeted against glycogen synthase 1 (Gys1, rabbit, 1:400, ab40810, Abcam), glial fibrillary acidic protein (GFAP, mouse, 1:500, BD556330, BD bioscience) and ionized calcium binding adaptor protein 1 (Iba1, goat, 1:350, ab5076, Abcam). Sections were then washed with PBS and incubated at room temperature for 2 h with secondary antibodies (ThermoFisher Scientific) diluted in blocking buffer: Alexa Fluor 488 donkey anti-mouse IgG (H+L) (1:500, Invitrogen A-21202), Alexa Fluor 488 donkey anti-goat IgG (H+L) (1:500, Invitrogen A-11055) and Alexa Fluor 594 donkey anti-rabbit IgG (H+L) (1:500, Invitrogen A-21207). After incubation with DAPI, sections were mounted using Aqua-Poly/Mount (Polysciences, Inc., US). Images of the different brain regions were taken on Zeiss LSM880 Airyscan confocal microscope at 40x magnification (zoom factor 0.6) with z-stack of 0.45 µm.

### Electrophysiology

Cortical slices were prepared from 3-12 month old male and female mice as previously described^42^ with some modifications. Mice were deeply anesthetized with a solution of xylazine (20mg/kg) and ketamine (150mg/kg) and transcardially perfused with ice-cold dissection buffer (in mM: 87 NaCl, 3 KCl, 1.25 NaH_2_PO_4_, 26 NaHCO_3_, 7 MgCl_2_, 0.5 CaCl_2_, 20 D-glucose, 75 sucrose, 1.3 ascorbic acid) and decapitated. Brains were transferred into ice-cold dissection buffer aerated with 95% O_2_-5% CO_2_. 400 μm-thick thalamocortical slices were cut on an angled block^51^ using a vibratome Leica VT 1200S, and transferred to an interface recording chamber (Harvard Instruments) and allowed to recover for 1 h in nominal “low activity” artificial cerebrospinal fluid (ACSF) at 32°C containing (in mM): 126 NaCl, 3 KCl, 1.25 NaH_2_PO_4_, 26 NaHCO_3_, 2 MgCl_2_, 2 CaCl_2_, and 25 D-glucose. Slices were then perfused with a “high activity” ACSF which contained (in mM): 126 NaCl, 5 KCl, 1.25 NaH_2_PO_4_, 26 NaHCO_3_, 1 MgCl_2_, 1 CaCl_2_, and 25 D-glucose. Slices remained in “high activity” ACSF for 30min and for recordings. Slices from 6- and 12-month old mice were exposed to Mg-deficient ACSF containing (in mM): 126 NaCl, 7 KCl, 1.25 NaH_2_PO_4_, 26 NaHCO_3_, and 25 D-glucose for 30 min before and during recordings. Spontaneous extracellular multiunit recordings were performed using 0.5 MΩ tungsten microelectrodes (FHC) placed in layer 4 of primary somatosensory cortex (5 minutes per slice). Recordings were amplified 10,000-fold, sampled at 2.5 kHz, and filtered on-line between 500 Hz and 3 kHz. All measurements were analyzed off-line using custom Labview software. For visualization and analysis of neuronal activity bursts, traces were offset to zero, rectified, and low-pass filtered with a 0.2 Hz cutoff frequency. Using these processed traces, the threshold for detection was set at 15-times the root mean square noise. A burst was defined if the amplitude of the signal remained above the threshold for at least 50 ms. The end of the burst was determined when the amplitude decreased below threshold for >600 ms. Two bursts occurring within 600 ms of one another were grouped as a single event. Event amplitude was defined based on the filtered/rectified traces. For power analysis calculated for a single recording, the same offset, rectification, and low-pass filtering were performed as described for event detection. The power spectrum of this processed signal was performed over the entire 30 seconds of each trace, and then averaged over all traces. The following frequency bands were examined: low delta (0.2-1 Hz), delta (1-4 Hz), theta (4-8 Hz), alpha (8-13 Hz), beta (13-30 Hz), low gamma (30-59 Hz) and high gamma (61-100 Hz). Power in each frequency band was normalized to total power (0.1-100 Hz).

### Statistics

Data were graphed and analyzed with Prism Graphpad 9, depicting mean + standard error of the mean. Lomb-scargle periodograms (Matlab) were applied to calculate the power and peaks of ultradian oscillations in activity. Behavioral endpoints were compared using two-tailed, unpaired student’s T (two groups) or one-way ANOVA (three groups). UP-state measures (Fig. 6) were compared using the Mann-Whitney U test^42^. *, **, ***, **** depict p<0.05, <0.01, <0.001 or <0.0001 respectively.

## Results

LB accumulation and evidence of CNS neuroinflammation were appraised qualitatively in WT and KO mice at ∼6 weeks, ∼6-7 months (Fig. S1,2) and ∼1 year of age through PAS-D staining and immunohistochemical assessments of glycogen synthase 1 (GS1, which accumulate within LBs), GFAP and Iba1 expression (Fig. 1A). LBs, GFAP and Iba1 induction were prominently seen in several brain regions, including the hippocampus, piriform cortex, striatum, cerebellum and the cochlear nucleus (Fig.S1-2). At 1 year age, KO and WT littermate mice were of similar body weight and responded similarly during an initial 2-hour long introduction to home-cage chambers (Fig. 1B-D). Within this highly enriched open field environment, we observed no significant differences in kinematic measures of home-cage exploration (obtained via videotracking) as well as objective measures of spout/feeder engagement (measured through lickometers and infrared beam-breaks, respectively). Even within this initial trial, both genotypes displayed a clear preference for sucrose containing fluid..

**Figure 1.**
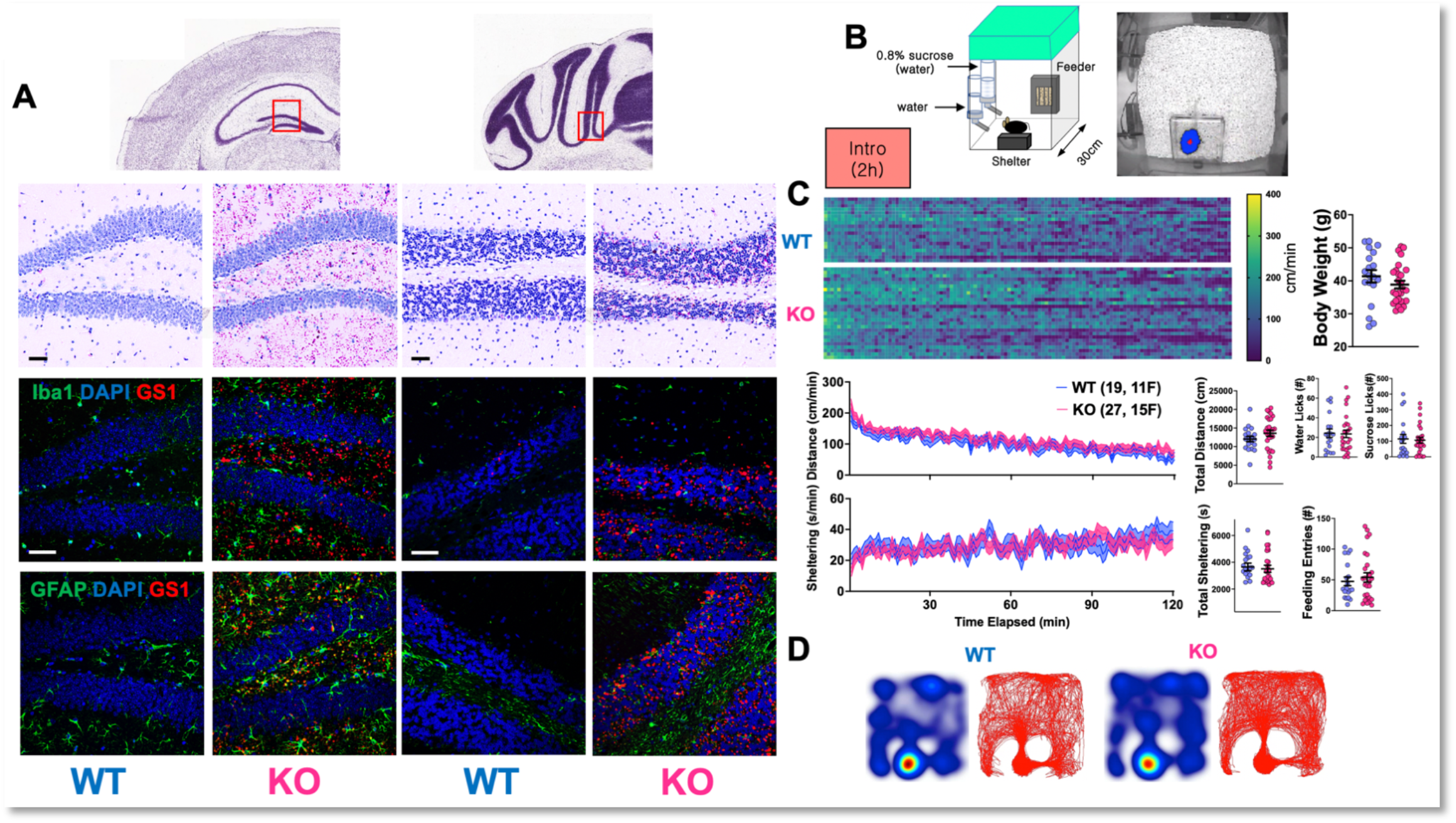
Lafora body (LB) Accumulation and Initial Home-Cage Response. A: Representative PAS-D images showing LB accumulation (TOP) and immunohistochemical assessments of LB accumulation (GS1), astrogliosis (GFAP) and microglial activation (Iba1). Scale bar: 50 µm. B: Home-cage structure and representative aerial video snapshot. C: Raster plot of distances traversed every minute of the 2-hour long introduction trial for every mouse, with measures of licking, shelter and feeder engagement. D: Heatmaps (left) and trackmaps (right) for a representative WT and KO mouse. Mean + s.e.m shown for all

During the second of two prolonged “baseline” recordings (allowing for a habituation day), WT and KO showed comparable patterns of nocturnality, total and hourly horizontal displacement, a similar mid-active phase dip in movement and nearly identical ultradian rhythms of activity (Fig. 2A). The timing and total amounts of “sleep” and “sleep bouts” (assessed noninvasively) were also equivalent (Fig. 2B). Over the entire trial, averaged time budgets^30,34-36^ allotted to sheltering, feeding and licking behavior were similar (Fig. 2C), and both groups of mice demonstrated a similar micro/macrostructure of licking/feeding bouts (Fig. 2D, E). Together, these data illustrate that within a relatively task-free home-cage situation, 1-year-old WT and KO mice display indistinguishable patterns of spontaneous behavior.

**Figure 2.**
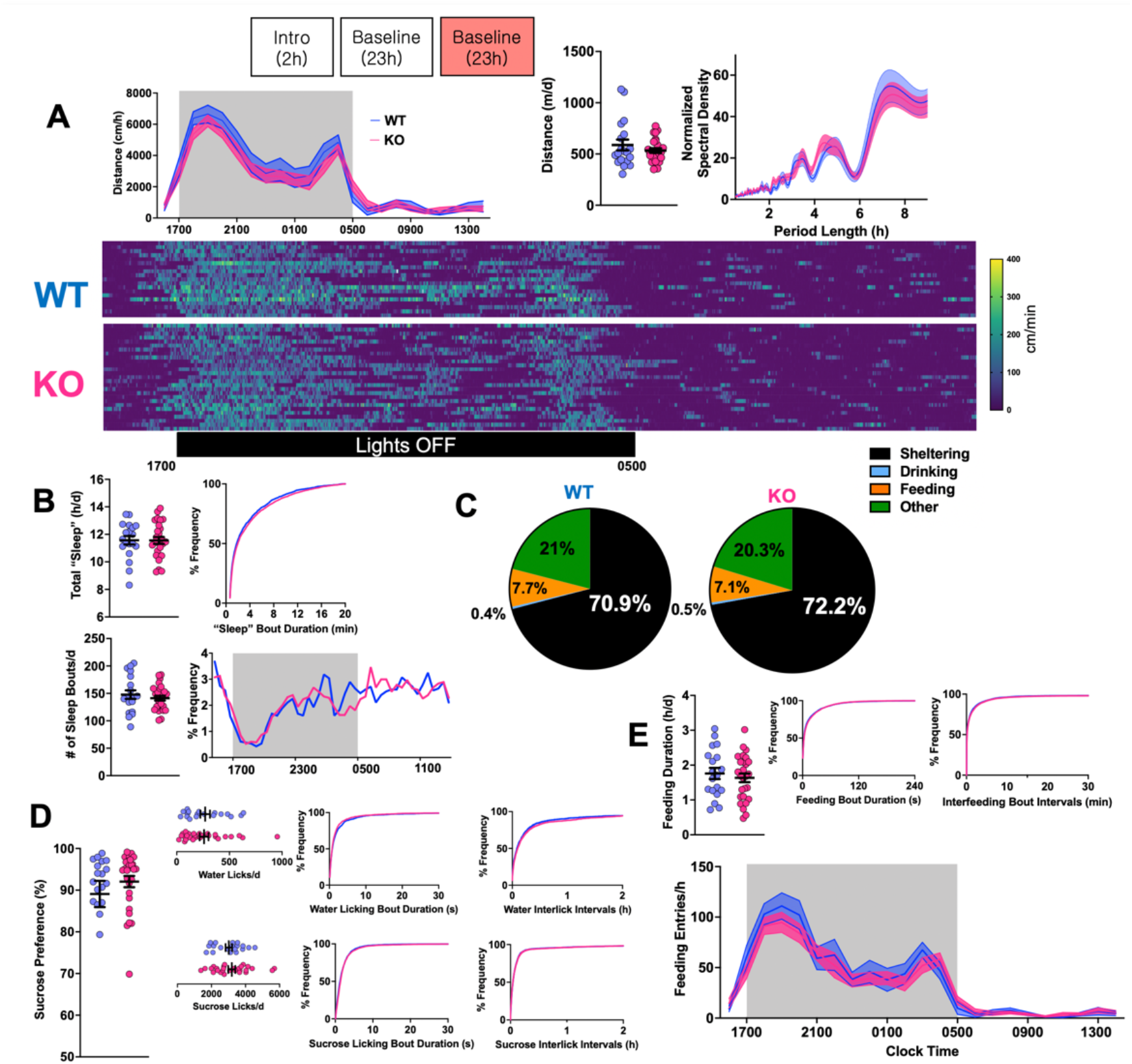
Baseline Recordings. A: Hourly horizontal distances within the home-cage (lights OFF between 1700 and 0500), total distances, and ultradian rhythms of activity (Lomb-Scargle Periodo-gram). BOTTOM: Raster plot of distances moved every minute of the day, displaying active/inactive state structure. B: KO mice displayed similar bout structure and timing of “sleep” (measured noninvasively). C: Averaged time budgets for WT and KO mice. D: WT and KO mice displayed similar sucrose preference and lick macrostructure. E: WT and KO mice displayed similar feeding entries and durations. Mean + s.e.m shown for all. See Fig.1A for sample sizes.

Having now profiled their spontaneous behavior within this task-less setting, we next presented a set of provocative maneuvers to reveal other latent phenotypes. During a light spot test^34-36,47^, a bright ceiling-mounted home-cage LED was illuminated for 60 minutes in the early nocturnal period (1900). WT and KO mice displayed equivalent responses (Fig. 3A). In comparison, KO responses to a 60-s long auditory tone (“beep”) were relatively blunted, with a weakened startle response and less vigorous shelter engagement (Fig. 3B). When presented with running wheel access, total wheel rotations were not significantly different (Fig. 3C). Finally, during a “swap” protocol^36^ where every mouse is repositioned within a cage previously inhabited by a sex-matched mouse, WT and KO mice displayed similar responses to this geometrically similar (yet olfactorily distinct) environment (Fig. 3D). Home-cage behavioral data (obtained through an identical modular design) at 6-week and 6-month timepoints are shown in Fig. S3 and S4 respectively. With the exception of wheel-running behavior at the 6-month timepoint (significantly lower in KO mice, p<0.05), no significant changes were observed across a large set of core behavioral parameters.

**Figure 3.**
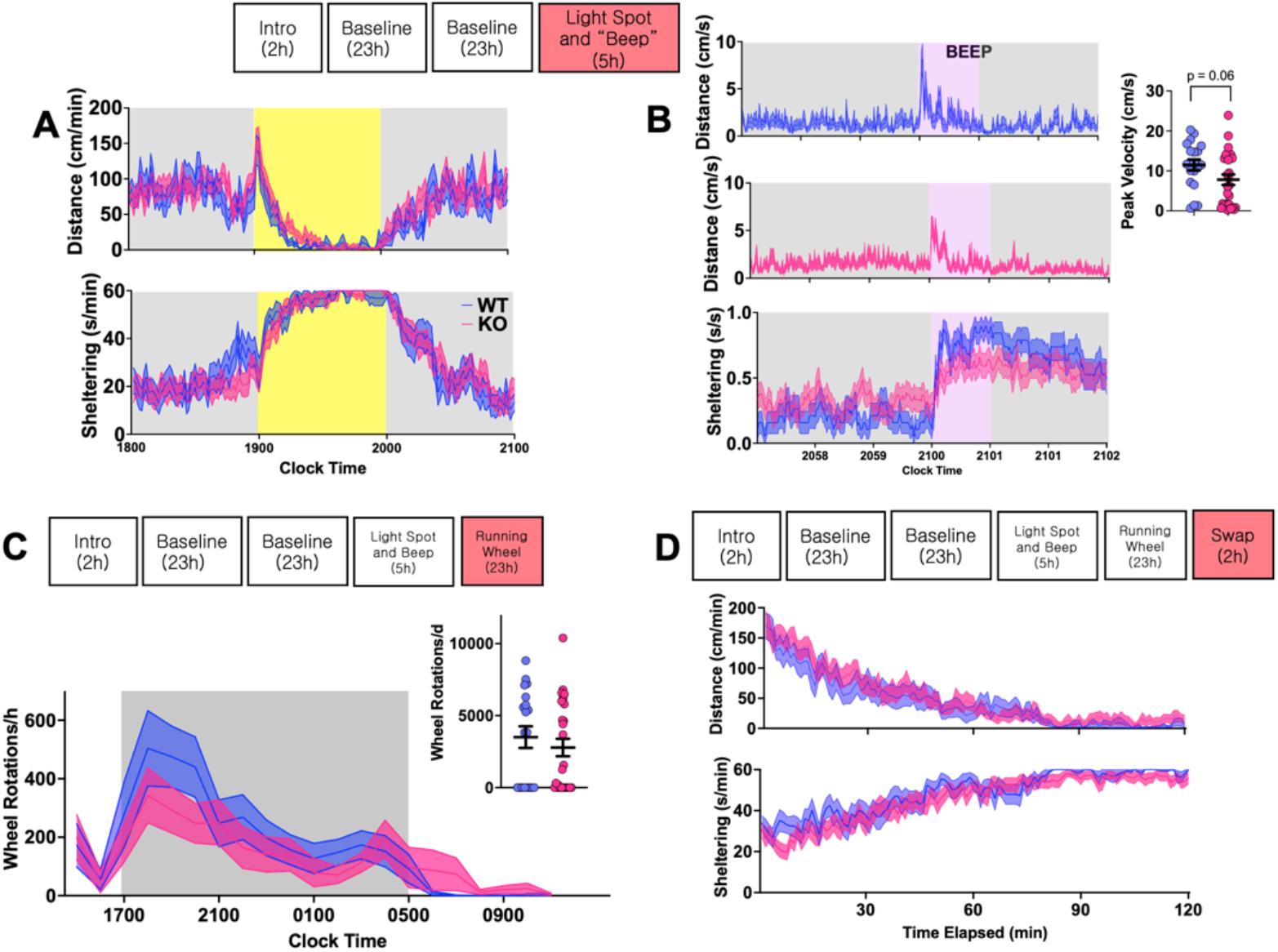
Home-cage provocative maneuvers. A: Changes to horizontal activity and sheltering in response to a single hour of light-spot stimulation. B: Distances and sheltering responses to a single 60s-long beep stimulus (2300hz tone). C: Wheel rotations accumulated during a 23h-long wheel-running trial. D: Distance/sheltering behavior following cage-swap. Mean + s.e.m shown for all. See Fig.1 for sample sizes.

Next, to explore whether any of our home-cage metrics varied by age, we directly compared the performance of 6 weeks, 6 month and 1 year old WT mice, acknowledging two main caveats to such a comparison. First, WT mice of differing ages were not studied simultaneously (we emphasized synchronous assessments KO and their WT littermates). Second, to reduce^41^ the number of animals employed, 13 WT mice were studied serially across all three timepoints. Compared to 6-week-old mice, those aged 6-7 months and older displayed a significant reduction in total feeding durations (but not feeding entries) and wheel-running interest (Fig. 4A, E). By one year of age, we observed a mild statistically significant reduction in sucrose preference. Total horizontal distances were also considerably lower at this timepoint, replicating previous results^38,52^ (∼587m/d, compared with ∼726m/d [6 weeks], 701 m/d [6 months]), although this effect was not statistically significant (p = 0.08). Estimates of total sleep time, timing and sleep structure remained largely unchanged (Fig. 4B). Compared with 6-week-old mice, older cohorts displayed more sustained shelter engagement during the light spot test (Fig. 4C) and a less pronounced initial startle response to auditory stimulation (Fig. 4D).

**Figure 4.**
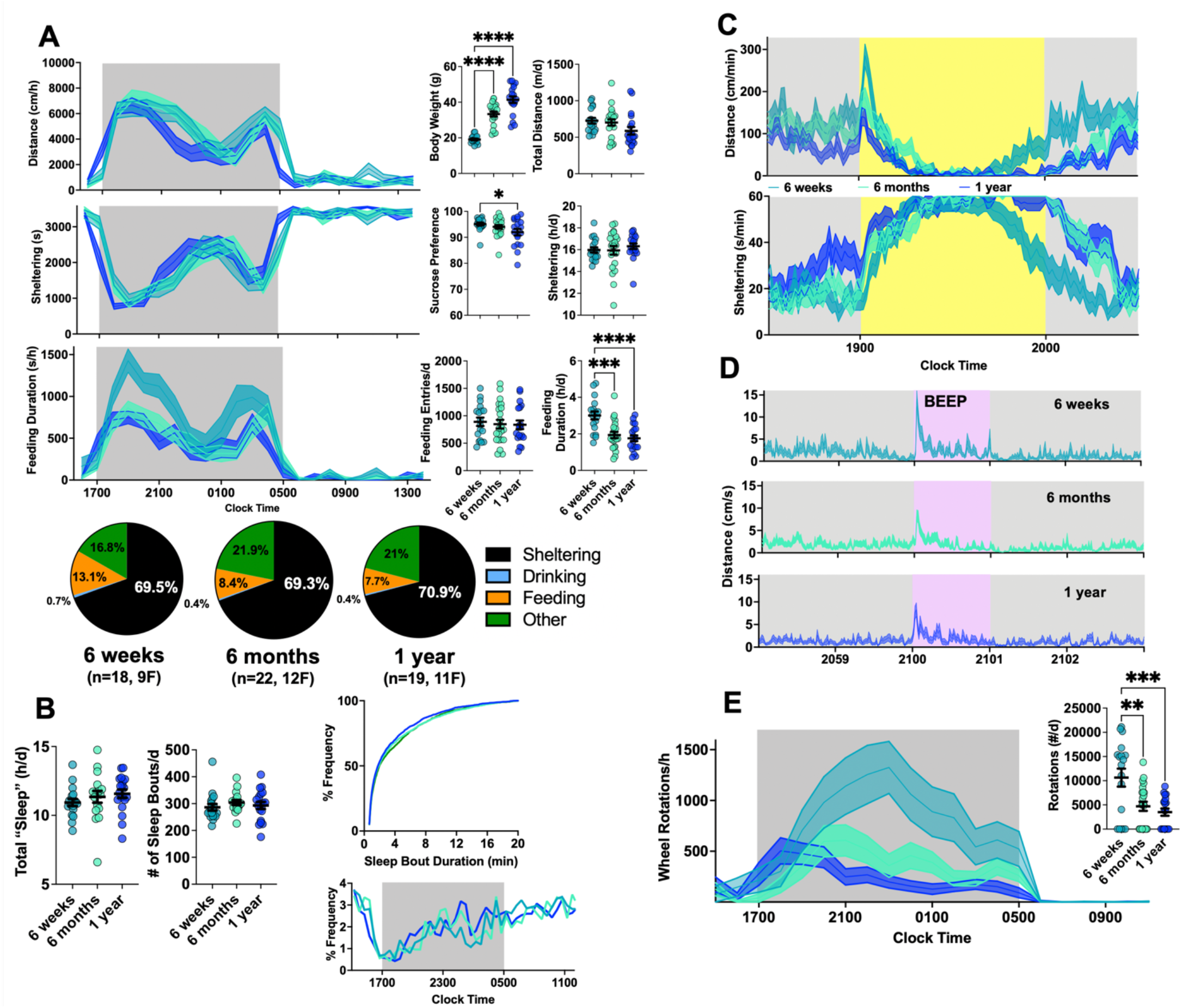
Changes in Home-cage Behavior with Age. A: On baseline day 2, compared with 6-week old mice, older WT cohorts displayed diminished sucrose preference (F_2,56_ = 2.94, p = 0.06) and feeding durations (F_2,56_ = 10.55, p<0,0001). B: Measures of sleep timing and duration were unchanged. C,D: 6-month and 1-year old mice displayed a blunted response to light-spot and BEEP stimulation. E: Older mice displayed fewer wheel rotations (F2,56 = 9.10, p<0.001). Mean + s.e.m shown for all. *, **, ***, **** depict p<0.05, <0.01, <0.001 or <0.0001 respectively

To survey for spontaneously occurring epileptic seizures at 1 year of age, a total of 4 WT and 7 KO mice underwent implantation of wireless EEG electrodes^36^, with each mouse receiving at least 96 hours of single channel EEG recording (Fig. 5A). Spontaneous seizures were not observed across either genotype. EEG spectral signatures (assessed during waking periods) were similar, featuring a peak in power at ∼4Hz (Fig. 5B). To ensure that our platform was sensitive to seizure activity, a subset of WT and KO mice received intraperitoneal injections of PTZ at convulsant doses (60mg/kg). As shown in Fig. 5C, PTZ-induced electrographic seizure activity was similar between WT and KO mice, beginning with quasiperiodic discharges (0.1-0.2Hz), progressing ultimately to an epoch of evolving rhythmicity, followed by post-ictal amplitude suppression.

**Figure 5.**
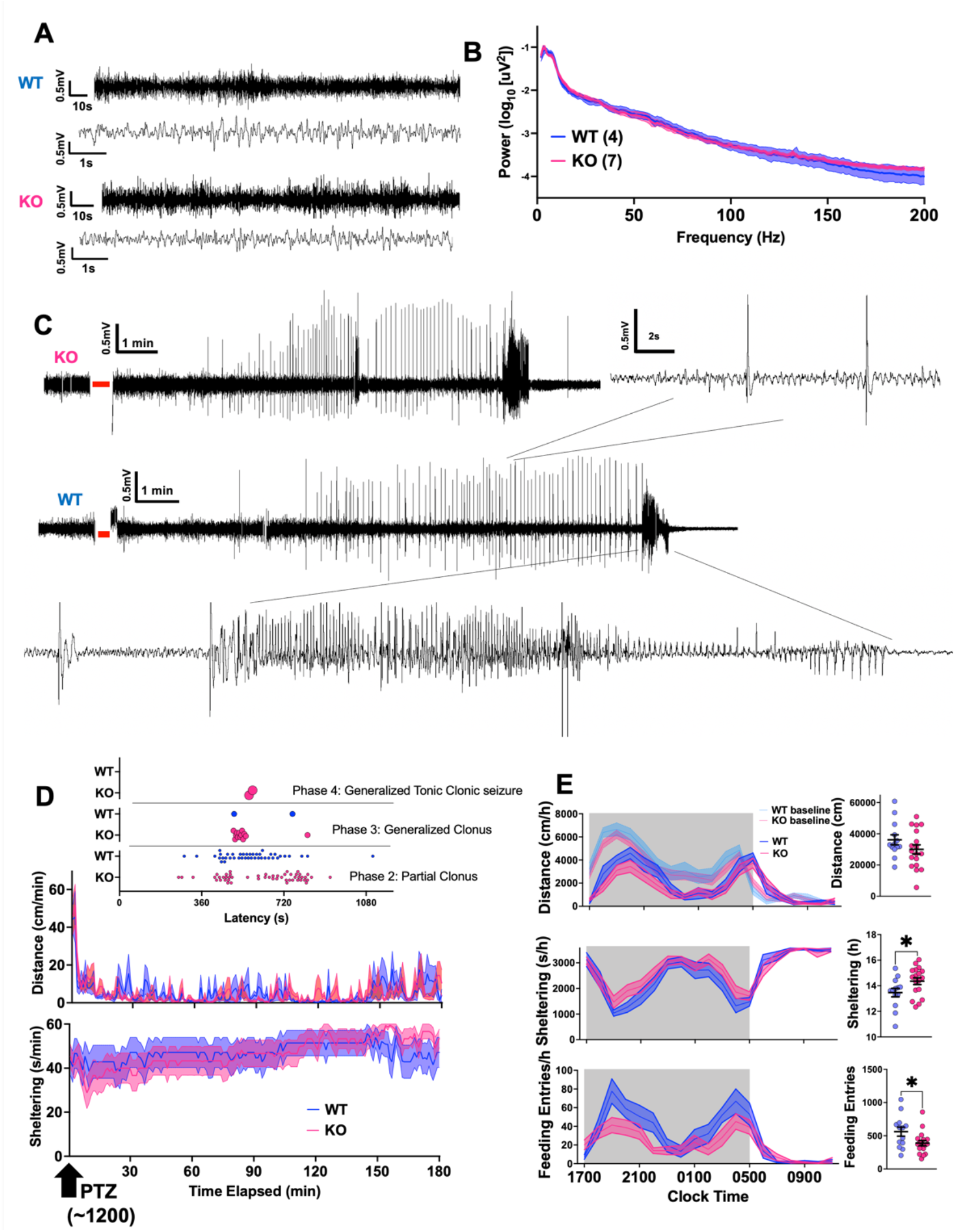
EEG and PTZ Responses. A: Representative single-channel electrocorticography from 1-year old WT and KO mice. B: EEG power spectra calculated during wakefulness. C: Representative EEG responses to a single intraperitoneal injection of PTZ (60mg/kg), demonstrating a prolonged epoch of spike/wave discharges, followed by a discrete epoch of evolving rhythmicity. Red bars annotate epochs of absent EEG signal while the mouse receives the intraperitoneal injection. D: Distance and sheltering responses to a single subconvulsant PTZ injection (30mg/kg), with a tally of convulsive events (inset) for both WT (n=14) and KO (n = 18). E: Post-ictal period home-cage metrics, revealing a comparative increase in sheltering and reduction in feeding in KO mice. Mean + s.e.m shown for all. * depicts p<0.05.

**Figure 6:**
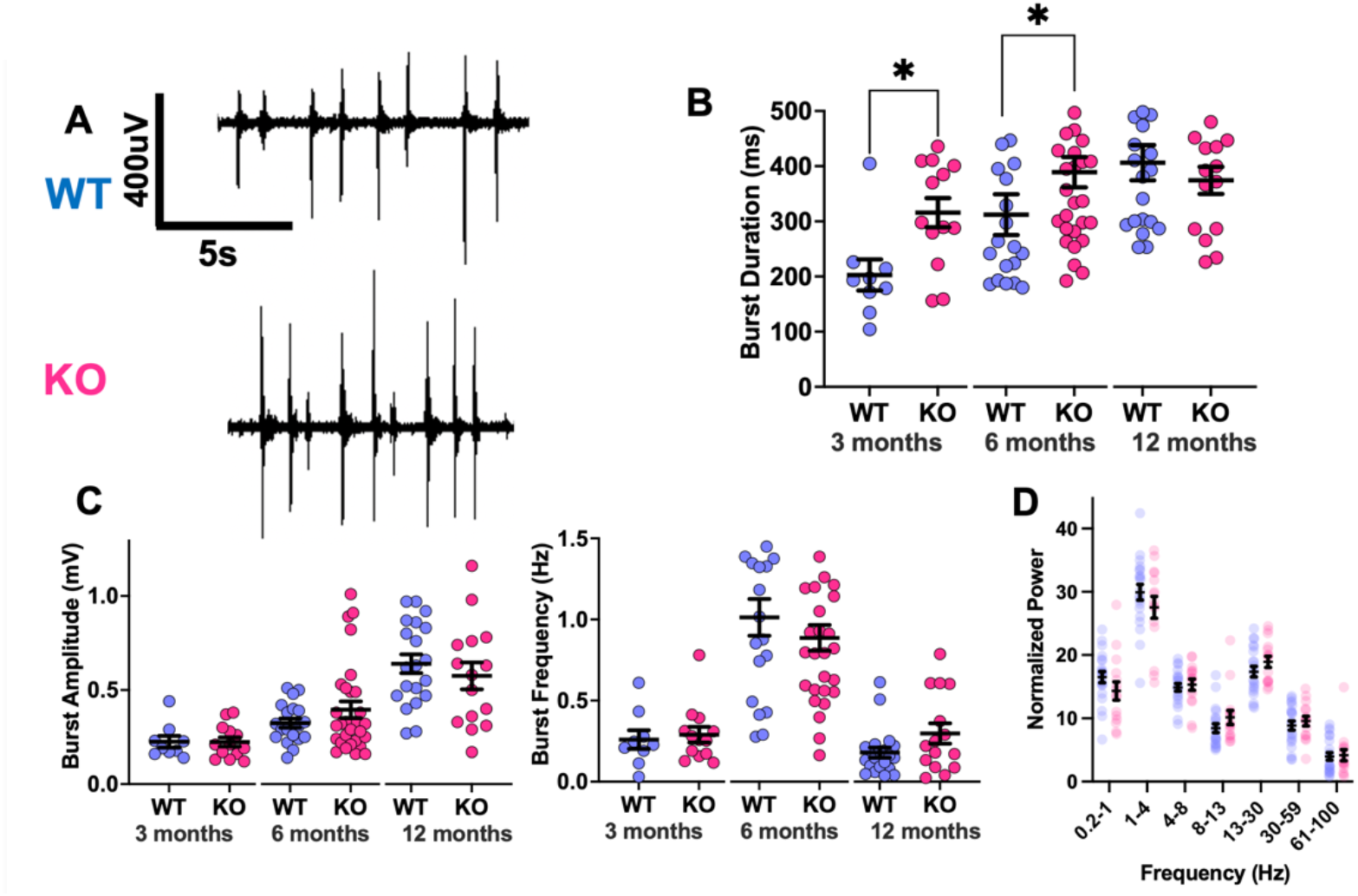
UP States in WT vs KO Somatosensory Cortex. A: Example traces of extracellular recordings in brain slices exhibiting spontaneously occurring activity bursts (from 12-month-old mice). B: The average duration of activity bursts in MKO slices is longer at 3 and 6 months of age, but not at 12 months (Mann-Whitney test, *p< 0.05) C: Burst amplitudes and frequencies. D: Relative power over all activity. Mean + s.e.m shown for all. Sample sizes at 3 months: WT (9 slices, 4 mice), KO (13 slices, 5 mice). 6 months: WT (19 slices, 6 mice), KO (29 slices, 7 mice). 12 months: WT (12 slices, 6 mice), KO (15 slices, 5 mice).

We then took advantage of our home-cage monitoring platform to objectively compare WT and KO behavioral responses to PTZ. Using videotracking data, we have previously conducted a detailed quantification of the remarkable immobility that is observed following a single intraperitoneal injection of subconvulsant dose PTZ (30mg/kg, administered at ∼1200). Since much of this early immobility occurs *outside* the shelter at a time of the day when mice are largely confined to their shelters^35^, this response quantifies the severity of the acute [ictal] encephalopathy induced by PTZ. As shown in Fig. 5A, changes in horizontal activity and shelter engagement were similar between genotypes. To tally the occurrences of clonic (phase 2, 3) and tonic-clonic seizures (phase 4)^48^, we manually examined video recordings and high-resolution actograms for each subject during the first 20 minutes following the PTZ injection (supplemental movie 1-3). KO mice displayed a greater incidence of generalized clonus and were the only genotype to display generalized tonic-clonic seizures. To recognize genotypic differences during a more extended post-ictal period, we profiled behavioral patterns over the remainder of the day. Between 1600-1100, KO mice displayed significantly greater shelter engagement and fewer feeding entries. While both genotypes displayed impressive post-ictal hypoactivity, this was similar between WT and KO mice (Fig. 5E).

Finally, in a separate group of WT and KO mice (without prior PTZ or other experiences), we asked whether LB accumulation and associated neuroinflammatory changes were associated with *in vitro* evidence of neurophysiological evidence of cortical circuit dysfunction. We adopted an approach that utilizes neocortical slices to record the expression of UP states^42,53,54^, which are states of synchronous depolarization driven by local recurrent excitation and inhibition within all neurons in a cortical region^55^. While UP states can be evoked with thalamic stimulation^56^, spontaneously occurring UP states are thought to underlie neocortical slow oscillations during slow wave sleep^57^. Spontaneous UP states are more prolonged in mice with deletions of *Fmr1* (modeling fragile-X syndrome^42^), while significantly shorter and less frequent UP states are seen in a mouse model of Down syndrome^58^. In our experiments, we found that UP state durations were significantly larger in KO mice at 3 and 6 months of age, but we observed no differences at 12 months of age. UP states were of higher amplitude^54^ and longer duration in older mice.

## Discussion

In this study, we examined one^16^ out of four^14,15,17^ available mouse models of malin-deficient Lafora disease to ask whether brain LB accumulation and associated astrogliosis/microglial activation are associated with any robust alterations in home-cage behavior. The home-cage approach affords several benefits that improve rigor and reproducibility of behavioral phenotyping^29,41^. This includes the automated acquisition of prolonged recordings in an experimenter-free setting, providing a particularly transparent window into the rich expressions of spontaneous behavior during the murine night. To avoid the potential observer effects associated with prolonged social isolation, we applied a modular design over a 4-5 day observation period designed to gauge behavioral wellbeing across multiple dimensions, including rest/arousal systems (distances, “sleep”), rhythms of consumptive behavior, responses to potential threats (e.g., light spot, swap) and reward responsiveness (e.g., wheel-running, sucrose preference). While abundant LBs were evident at 1 year of age (Fig. 1A, 7), KO mice were largely similar to WT mice across a range of scalar home-cage endpoints. As an internal control, we *did* identify important age-related changes in many of the same metrics within WT mice, including with older mice displaying lower sucrose preference, feeder engagement and wheel-running drive. These findings, in conjunction with previously published home-cage phenotypic distinctions across inbred strains of mice^59,60^ and disease models^29,34-36,38-40^ studied using the same apparatus, argue against platform insensitivity as an explanation for our negative findings.

**Figure 7:**
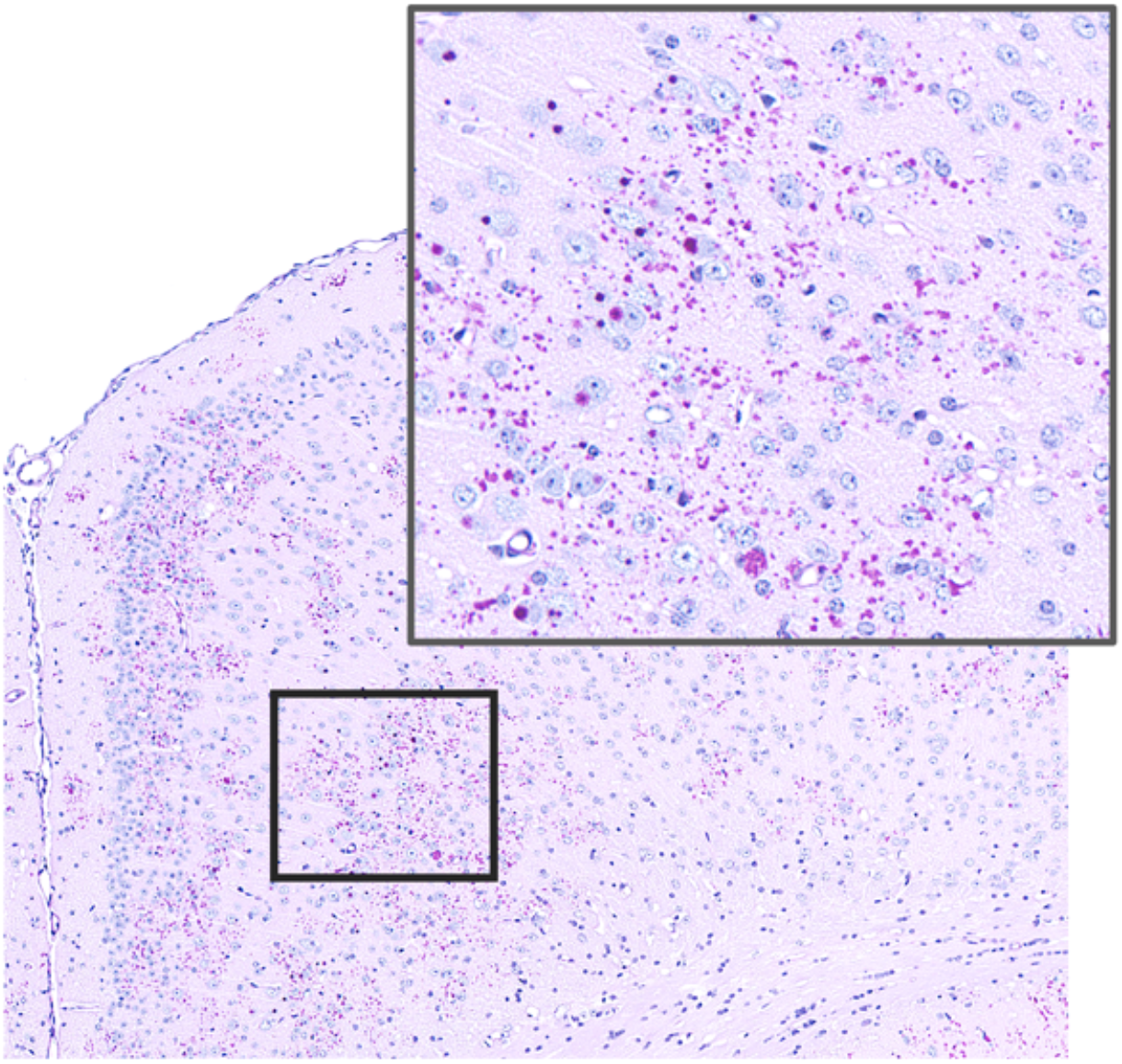
Lafora body accumulation in mouse piriform cortex (malin KO, 12 months of age).

In a slice preparation, we did find evidence for LB-associated neocortical dysfunction at 3 and 6 months of age, where UP state bursts were significantly prolonged without changes in burst frequency or amplitude. This may reflect a unique window of time for follow up studies designed to test the effects of LB scavenging or other therapies on neocortical dysfunction. However, differences in burst duration were not identified at 12 months of age. In alignment with these findings, EEG recordings did not identify significant changes in spectral composition between WT and KO mice at this time point. Nevertheless, in response to PTZ, KO mice displayed evidence of greater convulsive and behavioral seizure severity, reproducing earlier results^20,21^. While PTZ seizure induction paradigms can be quite heterogeneous across laboratories^61^, these data suggest that detailed analyses of PTZ responses may serve as a potential readout of cortical function.

We posit three possible non-mutually exclusive explanations for the clinicopathological dissociation that we uncover. First, laboratory mice may display a species-specific immunity to the neurobehavioral consequences of abundant LB accumulation. Second, clinically meaningful changes in neuronal dysfunction may substantially lag behind neuropathological abnormalities at a time scale (years?) that cannot be practically explored in laboratory mice. This explanation is compatible with LD in human and canine subjects^62-64^ (for whom serial neuropathological assessments are impossible), where epilepsy and neurocognitive decline occur after years of seemingly normal brain development. Third, there remains the possibility that LB accumulation, while impressive, is not the proximate cause of neuronal dysfunction related to the loss of malin, whose functions beyond glycogen metabolism remain unknown. We regard this as an unlikely possibility, since recent work has shown that malin’s predominant (if not exclusive) subcellular localization is at glycogen, where it is tightly scaffolded to the carbohydrate binding domain of laforin^11^, the deletion of which produces an identical clinical syndrome. In a number of LD mouse models (including the one studied here), preventing LB formation through downregulation of glycogen synthesis or removing LB by digesting them with a CNS-delivered amylase, prevents or corrects the neuroinflammatory, neurometabolic and brain protein glycation defects that characterize the neuropathology of LD^19,28,65-70^. However, the neurobehavioral correlates of these biochemical rescue strategies have not been explored in as great detail.

We identify three main limitations to this work. (i) Our neurobehavioral survey did not include any classical measures of learning and memory. As in humans, this is a multilayered construct in mice, with an array of available tests that are designed to assay fear memory (e.g., fear conditioning), spatial memory (e.g., Morris water maze), object memory (e.g., object recognition testing) or procedural memory (e.g., illuminated radial arm maze). Our results suggest that any learning/memory phenotypes, if present, cannot be explained by (or associate with) concurrent deficits in sleep, motor function, or grossly assayed visual or auditory function. (ii) Our insights could have been strengthened further by comparing two or more mouse models of LD, so as to more definitively correlate LB accumulation with home-cage behavioral deficits. (iii) Extending our recordings to even older mice (e.g., 21-27 months of age^38^) may have revealed additional phenotypes. (iv) And finally, as with any several-day long EEG survey in mice, the absence of spontaneous seizures does not necessarily imply seizure freedom per se, as KO mice may display extremely rare seizures that require longer EEG surveillance. Spontaneous seizure occurrence in mice is frequently associated with seemingly unexplained premature mortality^71^, which was not seen in our KO mice.

In conclusion, given the urgent need for LD treatments, it remains reasonable to screen for therapies that impart biochemical improvements in LB burden and related neuropathological changes. Our results find little evidence for robust changes in home-cage behavior in malin-deficient mice aged to approximately half their typical laboratory lifespan. Identifying holistic neurobehavioral endpoints to practically validate a pipeline of disease-modifying LD treatments may require us to innovate strategies that go beyond the laboratory mouse^64^, and justifiably invest in protocols that involve prolonged trial durations (years) that more closely mirror human LD.

## Acknowledgements and Funding

We would like to thank UT Southwestern Medical Center Whole Brain Microscopy Facility (RRID: SCR_017949) and the Quantitative Light Microscopy Core, a Shared Resource of the Harold C. Simmons Cancer Center, supported in part by an NCI Cancer Center Support Grant, 1P30 CA142543-01 and 1S10 OD021684-01. We thank Kate Luby-Phelps for access to imaging equipment. This work was supported by NIH grants to VK (K08NS110924, R01NS131399), JRG/KMH (U54HD104461, R37NS114516) and BAM (P01NS097197).

## Data availability statement

Annotated raw data will be made available upon reasonable request.

## Patient consent statement

N/A

## Permission to reproduce material from other sources

N/A

## Clinical trial registration

N/A

## Supplementary Information

**Supplementary Figure 1.**
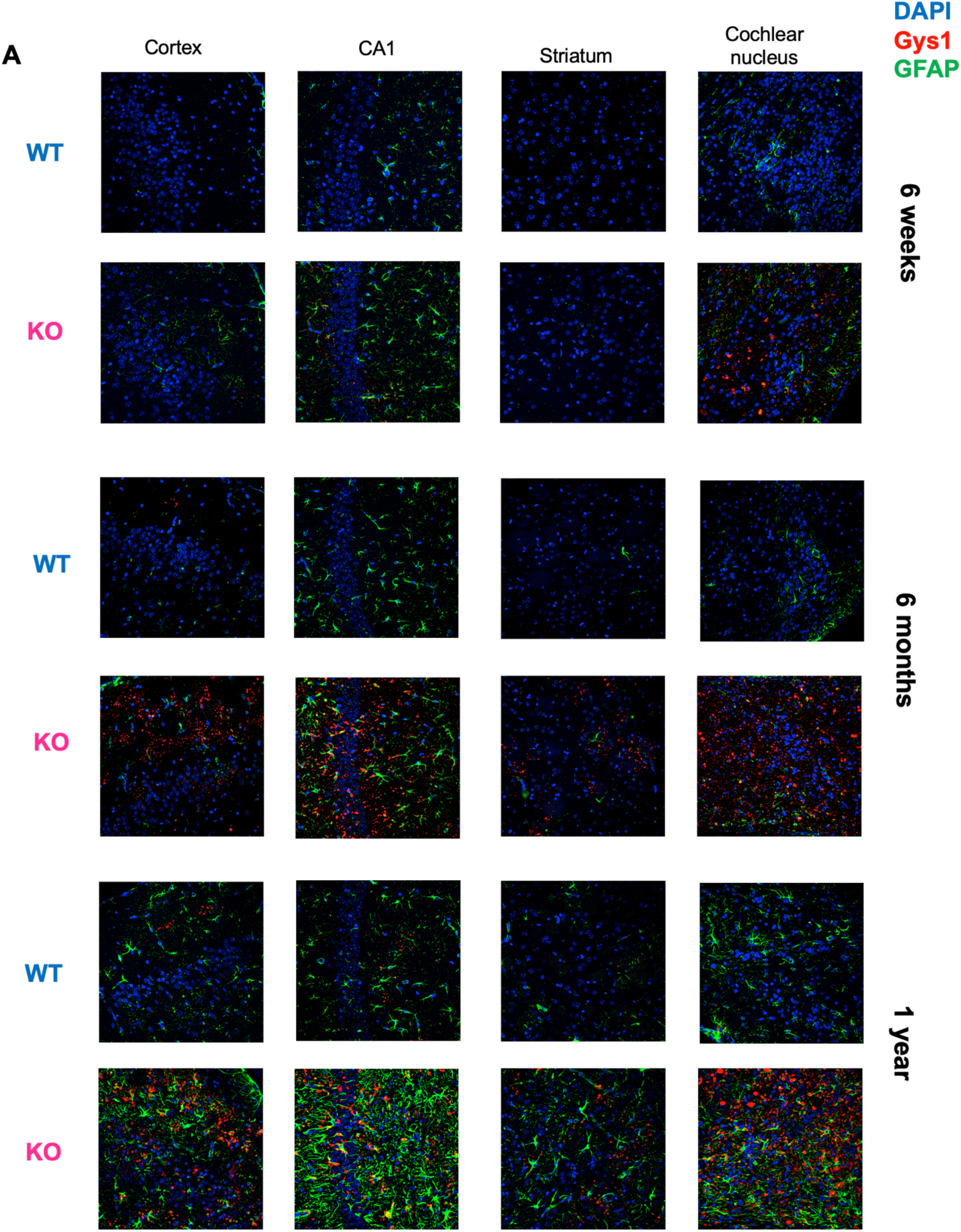
Age-dependent changes in Gys1 and GFAP expression (n = 2 mice/genotype)

**Supplementary Figure 2:**
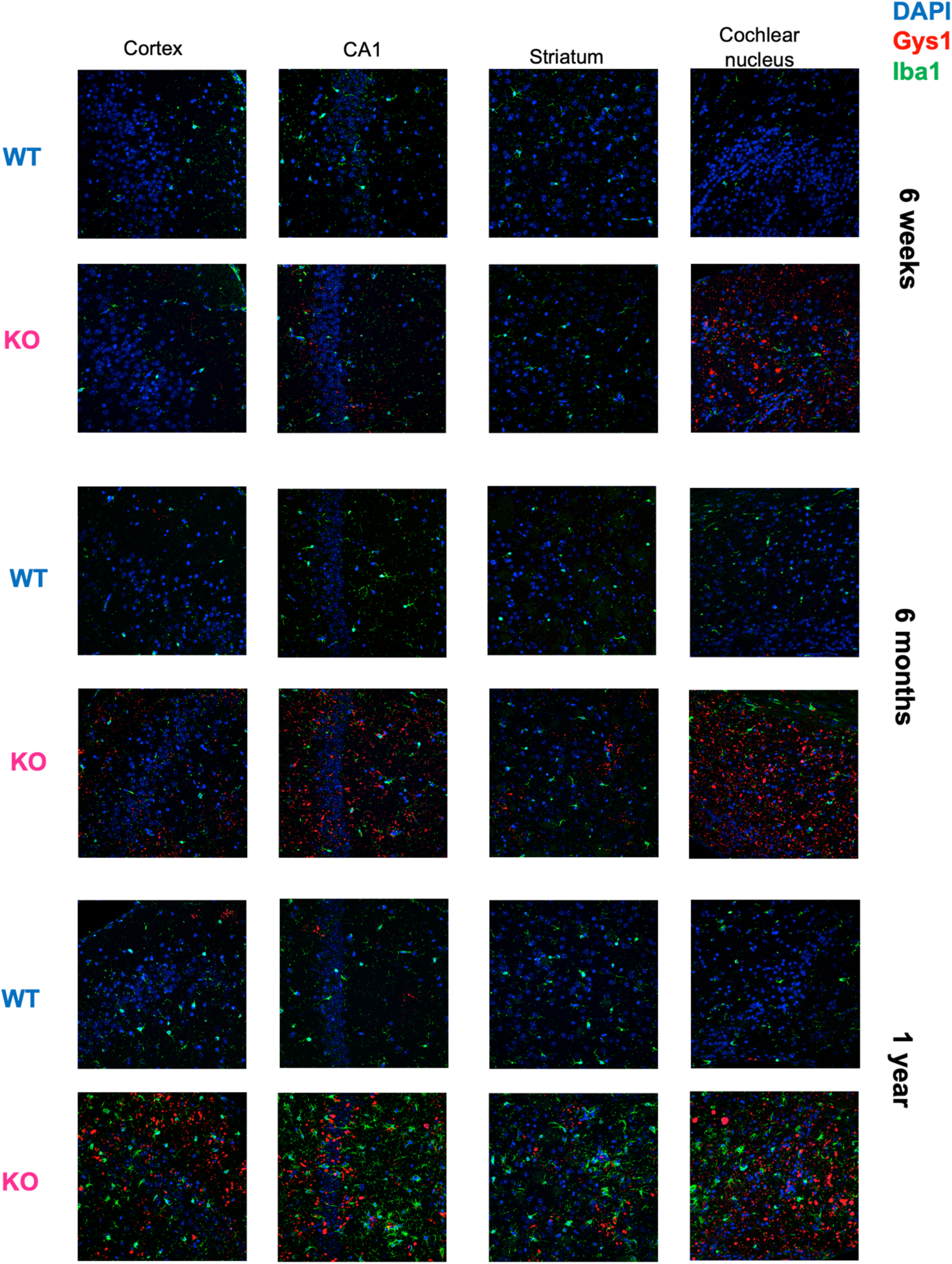
Age-dependent changes in Gys1 and Iba1 expression (n = 2-3 mice/genotype)

**Supplementary Figure 3.**
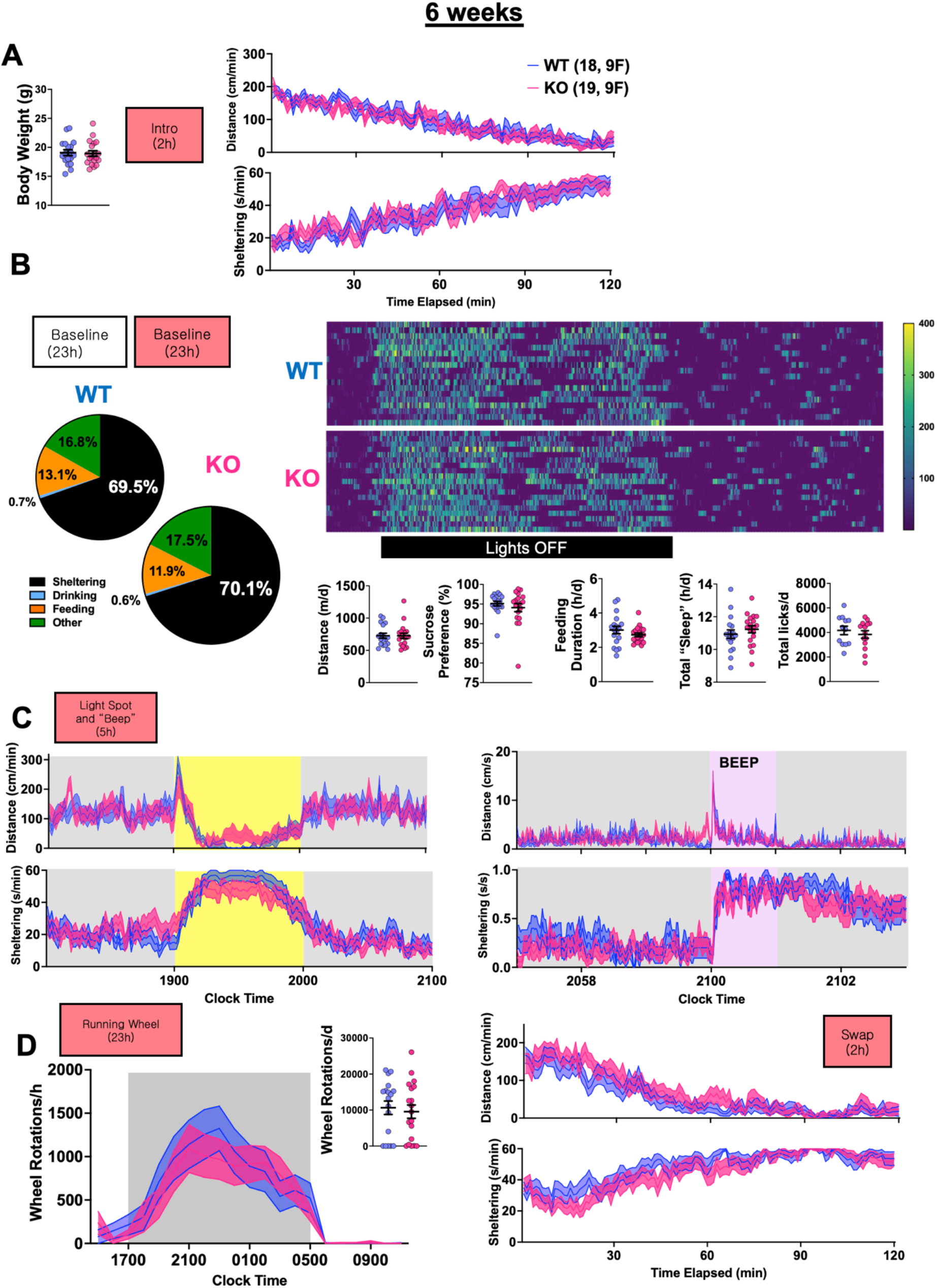
WT and KO Home-cage Behavior at 6 weeks of age. A: Initial body weights and exploratory responses when first being introduced to home-cages. B: Kinematic and consumptive behavioral metrics during the second baseline recording day. C: Responses to the light spot and BEEP stimuli within the home-cage. D: Wheel rotations (left) and responses to cage-swap (right). Mean + s.e.m shown for all.

**Supplementary Figure 4.**
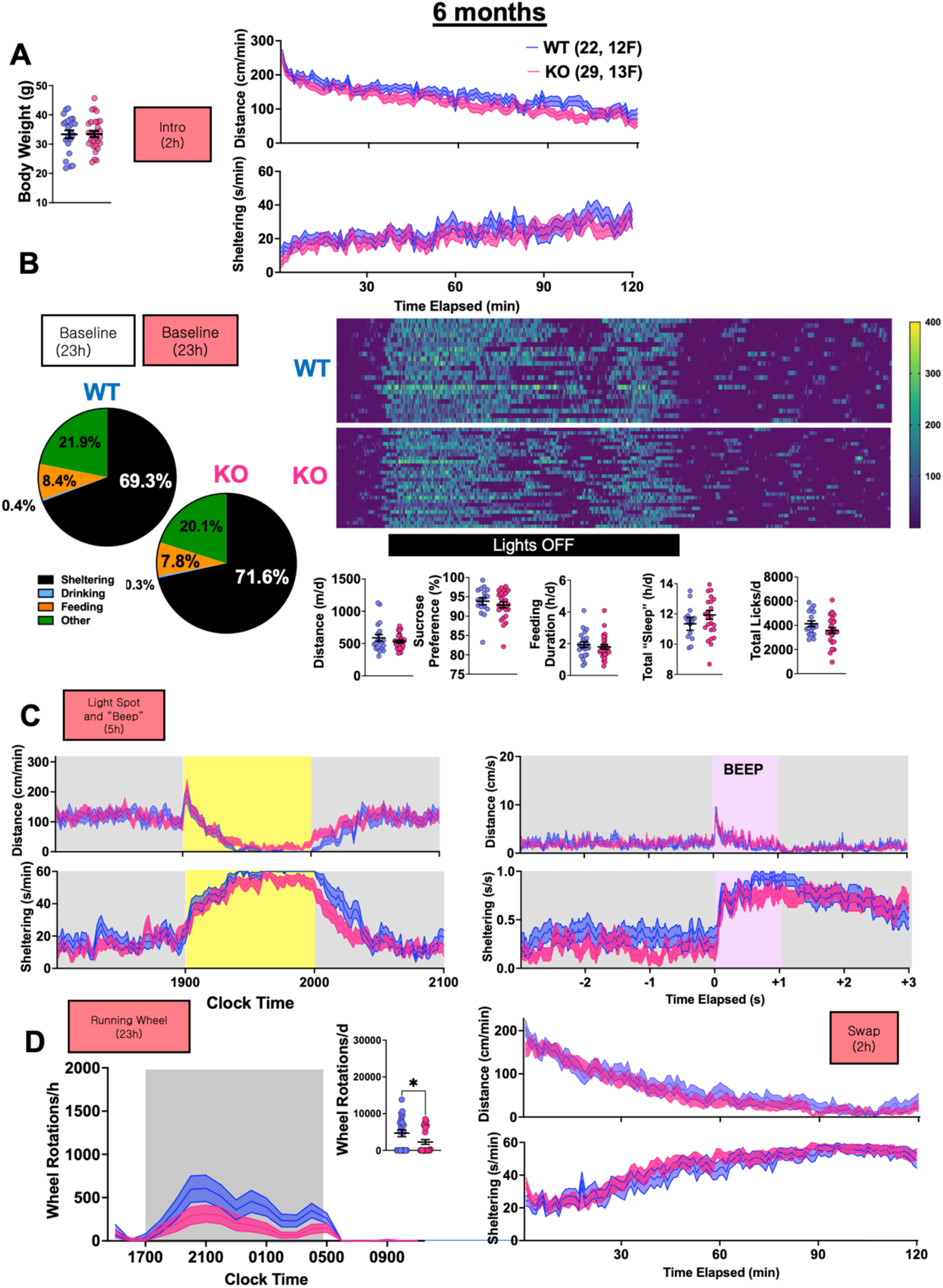
WT and KO Home-cage Behavior at 6 months of age. A: Initial body weights and exploratory responses when first being introduced to home-cages. B: Kinematic and consumptive behavioral metrics during the second baseline recording day. C: Responses to the light spot and BEEP stimuli within the home-cage. D: Wheel rotations (left) and responses to cage-swap (right). Mean + s.e.m shown for all. * depicts p<0.05.

**Supplementary Movie 1:** Representative video capturing two events scored as “phase 2” (4 and 8s into the recording), demonstrating clonic activity affecting the forelimbs.

**Supplementary Movie 2:** Representative video capturing a single “phase 2” followed by a “phase 3” event (7s into the recording), capturing generalized clonus manifesting as a sudden loss of upright posture.

**Supplementary Movie 3:** Representative video capturing a “phase 4” event (12s into the recording) featuring a *maximal* seizure without hindlimb extension.

